# Amd1-skp2 cascade regulates hepatocyte proliferation during liver growth and hepatocellular carcinoma development in zebrafish

**DOI:** 10.1101/2024.05.25.595867

**Authors:** Ke Zhang, Botong Li, Zhiling Deng, Yong Dong, Yuanyuan Li, Bingyu Chen, Ming Zhao, Mao Lu, Xingdong Liu, Zhenhua Guo, Sizhou Huang

**Affiliations:** Development and Regeneration Key Laboratory of Sichuan Province, Department of Anatomy and Histology and Embryology, School of Basic Medical Sciences, Chengdu Medical College, Chengdu 610500, China; Department of Neurology, the Second Affiliated Hospital of Chengdu Medical College, (China National Nuclear Corporation 416 Hospital), Chengdu 610000, China; Department of Gastroenterology, the First Affiliated Hospital of Chengdu Medical College, Chengdu, Sichuan, 610500, P.R. China; Department of Dermatovenereology, The First Affiliated Hospital of Chengdu Medical College, Chengdu, Sichuan, China; Ministry of Education Key Laboratory of Child Development and Disorders; Key Laboratory of Pediatrics in Chongqing, CSTC2009CA5002; Chongqing International Science and Technology Cooperation Center for Child Development and Disorders, Children’s Hospital of Chongqing Medical University, 400014, Chongqing, China; Central Hospital of Suining, Suining,629000, China

**Author notes:** **Correspondence:** (S. H); (Z. G).

**Keywords:** Zebrafish, Liver growth, Hepatocyte proliferation, *amd1*, *skp2*

## Abstract

Rapid hepatocyte proliferation is a common characteristic of liver growth in early liver development and hepatocellular carcinoma (HCC). Clarifying the mechanism how liver growth is regulated in development would benefit studying HCC progress. Although many genes have been reported to be involved in liver growth during early development, the mechanism underlying liver growth is far from being elucidated. Here we found that the liver undergoes rapid growth from 2dpf to 5dpf in zebrafish. Comparing the transcriptome for different staged hepatocytes identified 813 hepatocyte enriched genes at 3 dpf. Among them, S-adenosylmethionine decarboxylase proenzyme (AMD1) had not been previously reported to be involved in liver development. Our study further confirmed that *amd1* was highly enriched in hepatocytes during liver rapid growth and that *amd1* mutation inhibited liver growth mainly by repressing hepatocyte proliferation. Mechanistically, *amd1* loss of function downregulated the expression of *skp2*, and *skp2* is required for *amd1* to regulate hepatocyte proliferation. Furthermore, the role of Amd1-Skp2 cascade in regulating hepatocyte proliferation was also conserved in a zebrafish HCC model. In conclusion, our study systematically identified some uncharacterized genes possibly being involved in regulating liver growth during zebrafish development, and revealed the role of *amd1*-*skp2* cascade in regulating hepatocyte proliferation during liver growth and HCC progression. This work also provided a database which would benefit elucidating the mechanism how hepatocyte proliferation is regulated during liver growth and HCC progress.

## Introduction

The liver is one of the largest and critical internal organs that fulfils several critical functions, including metabolism, secretion, detoxification, and homeostasis (Li et al., 2021). Liver organogenesis involves multiple processes including ventral foregut endoderm migration, hepatoblast specification, liver budding, hepatoblast differentiation and the final step of liver growth and morphogenesis (Cheng et al., 2006; Field et al., 2003; Khaliq et al., 2015). In recent decades, the mechanisms by which liver development is regulated have been broadly studied. During this process, many factors, including fibroblast growth factors (FGFs), bone morphogenetic proteins (BMPs), Wnt signalling, hematopoietically expressed homeobox (Hhex) and (prospero-related homeodomain protein 1) Prox1, have been reported to be involved in hepatoblast specification and differentiation (Campbell et al., 2021; Goessling and Stainier, 2016; Tachmatzidi et al., 2021). In the later liver growth stage, *klf6* (Zhao et al., 2010), Hnf4α (Zhao et al., 2018), oestrogen (Chaturantabut et al., 2020), *hdac3* and Hippo signalling (Cox et al., 2016; Cox et al., 2018; Wu et al., 2022) have been reported to be specifically involved in hepatocyte proliferation (Ober and Lemaigre, 2018). Among all these factors, Yap seems to be most critical for hepatocyte proliferation and liver size control (Avruch et al., 2011; Wu et al., 2022). Although the mechanism underlying hepatocyte proliferation and liver growth has been broadly addressed, the detailed mechanism is far from being elucidated.

Recently, bulk RNA sequencing (RNA-seq) or single-cell RNA sequencing (scRNA-seq) have been used to study liver differentiation (Camp et al., 2017; Gao et al., 2022; Segal et al., 2019; Yang et al., 2017), regeneration (Ben-Moshe et al., 2022; Wang et al., 2019) and liver disease (Kimura et al., 2022; Saviano et al., 2020), but no study has screened genes highly expressed in hepatocytes during liver growth. To identify the genes involved in liver growth, we performed RNA-seq analysis to screen out liver-enriched genes at 3dpf using sorted GFP-labelled hepatocytes from embryos at 3dpf, 7dpf and adult zebrafish livers. Many novel genes, including S-adenosyl methionine decarboxylase proenzyme (AMD1), were identified, and their role in liver development had not been addressed. AMD1 is one of the key enzymes involved in the synthesis of polyamines (Bian et al., 2021; Pegg, 2009). During embryonic development and tissue regeneration, it is associated with embryonic stem cell (ESC) self-renewal, cell proliferation and cell migration (James et al., 2018; Lim et al., 2018; Zhang et al., 2012; Zhao et al., 2012). In pathological conditions, AMD1 is enriched in multiple cancers, including prostate cancer and neuroblastoma (Evageliou et al., 2016; Zabala-Letona et al., 2017), and AMD1 was also reported to be a potential target for tumour therapy since AMD1 blockade inhibits neuroblastoma progression. In hepatocellular carcinoma (HCC), AMD1 was enriched in human HCC tissues, and AMD1 increased HCC metastasis by stabilizing the interaction of IQGAP1 with FTO (Bian et al., 2021). Although the role of AMD1 in ESC self-renewal and tumour progression has been reported, whether AMD1 has a critical role in regulating liver growth has not been addressed.

Since *Amd1 ^−/−^* mouse embryos die between E3.5 and E6.5 days post-coitus (Nishimura et al., 2002), the role of AMD1 in liver development could not be evaluated in mouse embryos. To study the role of *amd1* in liver development during early embryogenesis, we generated two zebrafish *amd1* mutants using the CRISPR/Cas9 method. Our data showed that *amd1* loss of function inhibited liver growth. Mechanistically, *amd1* loss of function decreased the expression of S-phase kinase-associated protein 2 (*skp2*), a key component of the SKP1-cullin 1-F-box (SCF) complex (Wu et al., 2021; Zhang et al., 1995), and the downregulation of *skp2* in the *amd1* mutant enhanced the phenotype with decreased hepatocyte proliferation. In addition to the role of *amd1* in liver growth, our data also suggested that *amd1* was involved in HCC progression and that this role was also partially mediated by *skp2*. In conclusion, our data revealed some uncharacterized and liver-enriched genes during liver growth and further clarified the role of the *amd1-skp2* cascade in hepatocyte proliferation, including in liver development and hepatocellular carcinoma (HCC).

## Methods and materials

### Ethics statement

All experimental methods and protocols were approved by Chengdu Medical College (Sichuan, China). Zebrafish were maintained in accordance with the Guidelines of the Animal Care Committee of Chengdu Medical College.

### Fish and Fish Maintenance

Wild-type (AB), transgenic line *Tg(fabp10a:RTTA),Tg(Tetre:EGFP-kras_G12V)* (Nguyen et al., 2016)*, Tg(fabp10a:GFP)*, *skp2^+/-^ and amd1^+/-^* line fish were maintained in standard conditions at approximately 28.5 °C. The developmental stages were characterized as previously described (Kimmel et al., 1995).

### *Amd1* mutant construction

One sgRNA was designed according to the exon 1 region of the *amd1* gene, and the sgRNA was synthesized in vitro as described in the manual (HiScribe™ T7 High Yield RNA Synthesis Kit, NEB, NO. E2040S), the sgRNA and Cas9 protein were mixed and coinjected into the cell cytoplasm at the one-cell stage. Several injected embryos were used to evaluate the mutation efficiency using sequencing, and the remaining embryos were designated F0 (Founder). F0 adult fish was crossed with wild type to obtain F1 embryos. After evaluating the mutation efficiency using part of F1 embryos, the remaining F1 was maintained and grown to produce F2 embryos. Two types of useful mutations were identified in F2. The targeted sequence of *amd1 sgRNA1* was 5’-GGAGGTGTGGTTCTCCCGGC-3’. The primers used for amplifying the targeted genomic DNA (used for sequencing) were *amd1*-check*-*F: 5′-GTGATTCCATCCGACGGTTTA -3 ′ and*amd1*-check*-*R: 5′-CTGACATTATCACAGCGTTTCAC -3′.

### Bulk RNA Sequencing

To compare the transcriptome of hepatocytes in different liver stages, GFP-labelled hepatocytes were sorted using flowcytometry (Moflflo XDP, Beckman) from transgenic line *Tg(fabp10:GFP)* embryos and adult livers. Approximately 400 hepatocytes were collected for each stage. cDNA libraries were generated from these sorted cells using the Smart-seq2 protocol. RNA sequencing was performed using the PE100 strategy (HiSeq 2500, Illumina). Sequencing data were analysed as previously reported (Liu et al., 2016). To compare the transcriptome of wild-type and *amd1^7-/-^* embryos at 4 dpf, total RNA was prepared using TRIzol according tothe manual. RNA sequencing and analysis were performed by Novogene Co., Ltd. (Tian Jin, China).

### Fin-clip and identifying *amd1* mutation embryos

Since there is no clear morphological phenotype for *amd1^-/-^*embryos, to identify *amd1* homozygotes, the tail fin of zebrafish larvae was cut at 48 hpf, and genomic DNA was individually prepared as following: After anaesthetizing the embryo, the tip of the tail was cut with a scalpel to prepare genomic DNA as previously reported (Wilkinson et al., 2013), and the embryos were kept for further experiments. Genomic DNA was used to amplify the target region using PCR. Then, we evaluated whether the larvae were homozygotes according to the PCR results. For the wild-type larvae and heterozygotes, the target fragment was obtained, while for the homozygotes, the target fragment was not amplified, no amplicon was detected. The primers used here were as follows: *amd1*-screen-F: 5’-GTGGTTCTCCCGGCAG-3’, *amd1*-screen*-*R: 5′- CTGACATTATCACAGCGTTTCAC -3′.

### Cas9-sgRNA ribonucleoprotein complex (RNP) preparation and injection, *skp2* mutant generation

To obtain mosaic mutants in F0 embryos, we optimized the CRISPR-cas9 gene editing process according to a previous report (Wu et al., 2018). In brief, three sgRNAs for *skp2* were designed and synthesized in vitro using HiScribe™ T7 High Yield RNA Synthesis Kit(NEB, E2040S). The Cas9-sgRNA ribonucleoprotein complex (RNP) solution was prepared as follows: Cas9 protein (EnGen^®^ Spy Cas9 NLS, NEB, M0646T) 1.3μl, 1 M KCl 1.1μl, total sgRNA 2500 ng (830ng each), phenol red 0.3μl. Finally, RNA-free H_2_O was added to 5μl total, mixed completely, incubated at 37 °C for 5 minutes, and then placed back into an ice bath for the following injection procedure. The RNPs were injected into the embryo yolk within 15 minutes after fertilization. To examine the mutant efficiency for *skp2*, 16 injected embryos were used to prepare genomic DNA, and semi-quantitative RTLPCR was performed for evaluation as described in a previous report (Zhu et al., 2019). The primers used are shown in Table S1. To get the stable *skp2* mutant lines, the method being used to screen *amd1* mutant was used to screen *skp2* mutant. In our work a frameshift mutant line was obtained. The detailed information was provided in the result section.

### Chemical treatment

SMIP004 was used to inhibit the function of *skp2* as described in a previous report (Li et al., 2019). Two concentrations, 40 μM and 80 μM, were selected to inhibit *skp2* activity. The chemical SMIP004 was diluted with egg water to the concentration described above. The embryos were incubated with SMIP004 solution (40 μM or 80 μM) from 3 hpf to 24 hpf to evaluate which concentration was the proper concentration. Then, the embryos were incubated with SMIP004 solution (proper concentration: 80 μM) from 48 hpf to the stages needed.

### Plasmid Construction

Total RNA was extracted following the manufacturer’s instructions (TRIzol, Ambion, 15596-026). cDNA was prepared using a Revert Aid First Strand cDNA Synthesis Kit (Fermentas, K1622) according to the manufacturer’s instructions. The CDs of *amd1* and *skp2* were amplified individually using PCR (Prim STAR Max Premix Takara, R045A) and cloned into the PCS^2+^ vector (5x In-Fusion HD Enzemy Premix, Takara, 639649). The primers for cloning were as follows: PCS^2+^_F: 5′ -CTCGAGCCTCTAGAACTATAGTG-3 ′, PCS^2+^_R: 5 ′-TGGTGTTTTCAAAGCAACGATATCG-3 ′, *amd1-pcs2+_*F: 5 ′-TCTTTTTGCAGGATCGGAGTCTGTTTGTCTCACGATGG-3 ′, *amd1-pcs2+_*R: 5 ′-GTTCTAGAGGCTCGACGCTTCTTCATGTCAGAGGATCAG-3 ′, *skp2-pcs2+*F:5 ′- CTTTGAAAACACCACAAGTCAGGATGTCAAACGAAAGG-3 ′, *skp2-pcs2+*_R: 5 ′-GTTCTAGAGGCTCGAGCATTAATGTTTGTAGACGAGTCTGC-3′.

### mRNA injection

*skp2* mRNA was synthesized in vitro using an mMESSAGE Kit (Ambion, AM1340) as the described in the manual. The concentration for *skp2* mRNA injection was 40ng/μl. *skp2* mRNA was injected at the 1-4 cell stage.

### RT**□**qPCR

RTLqPCR was performed using the Brilliant III Ultra-Fast SYBR Green QPCR Master Mix (Agilent Technologies) and the CFX96 Real-Time System (BIO-RAD) according to the manufacturer’s instructions. The amount of *beta-actin* was used for normalizer. The primers are listed in Table S1. All experiments were repeated at least 3 times.

### Whole-mount *in situ* hybridization and section

One color *in situ* hybridization was performed as described in a previous study (Liu et al., 2019). The previous probes *fabp10, prox1, hhex,* and *fabp2* were used as described in previous reports (Zhang et al., 2022). The CDs of *amd1* and *skp2* were amplified using PCR and cloned into the vector pcs2^+^, then linearized the plasmids and synthesized the individual antisense probe as previously reported (Zhang et al., 2022). Two color *in situ* hybridization was performed as described in a previous study (Dunn et al., 2022). Specifically, digoxygenin-labeled *amd1* probe and fluorescein - labeled *uox* probe were used in our study. For sectioning, *in situ* hybridized embryos were re-fixed in 4% paraformaldehyde in PBS, followed by incubation in 15 and 30% sucrose in 0.1% Tween/phosphate buffered saline for 2 hours each. Then, embryos were mounted in 1.5% agarose in 30% sucrose, and balanced in 30% sucrose solution overnight at 4 °C. The mounted embryo was re-mounted in OCT (Sakura), sectioned using a CM1850 cryostat (Leica),

### Immunostaining

The embryos were fixed overnight with PFA (4% in PBS) at 4 °C, washed with PBS (5 min, 3x) and blocked with PBTN (4% BSA, 0.02%NaN_3,_ in PT) for 2 hours at 4 °C. Then, the primary antibody against H3p (GTX128116) or Caspase3 (BD 559565) was diluted with PBTN at 1:200 and incubated on a shaker at 4 °C overnight. Then, the embryos were washed with PT (0.3% Triton-X-100, in 1X PBS) for at least 20 min 8 times. The secondary antibody, Donkey anti rabit IgG, Texas Red coupled; GeneTex 26800) or Alexa FluorTM 647 (invitrogen, A21244) was diluted with PBTN in 1:500 and added. The embryos were incubated overnight at 4 °C (kept in the dark). Finally, the embryos were washed with PT more than 8 times (30 min each time) and imaged.

### EDU experiment

BeyoClick™ EdU Cell Proliferation Kit with Alexa Fluor 594 (Beyotime, C0078S) was used for this experiment. 200uM EDU solution containing 2%DMSO and 0.01% phenol red was prepared. Zebrafish embryos were mounted with low melting point agar (0.8%) and the EDU solution was injected pericardially. After injection, the embryos were replaced in egg water at 28.5℃ for 40 minutes, then fixed them overnight at 4L with PEM. The fixed embryos were rinsed with PBS for 3 times (5 min each time) and treated with pre-cooled acetone at −20 for 40 min, then were washed by PT for 3 times (20 minutes each time). Next the embryos were incubated with 3%BSA at room temperature for 2 hours and washed with PT 3 times, then according the manual the EDU reaction solution was added and keep the reaction for 30 minutes in dark. After EdU staining reaction the embryos were washed with PT for 3 times, flowing Immunostaining for GFP.

### TUNEL staining

Embryos were fixed in 4% PFA overnight at 4 °C, washed with PBST 3 times (10 minutes each time) and stored in 100% methanol overnight. Then, the embryos were washed 3 times with PBST 3 times, and an In Situ Cell Death Detection Fluorescein kit (Roche11684795910) was applied to examine cell apoptosis according to the manufacturer’s instructions.

### Microscopy

Images of whole-mount *in situ* hybridized embryos (mounted in 80%-100% glycerol) and section samples were captured at room temperature using an OLYMPUS SZX16. To examine positive pro-apoptotic, proliferating cells in the liver, the *Tg(fabp10:EGFP)* embryos were fixed in PFA (4% in PBS) overnight and mounted in 1.5% Low Melting-point Agar. Then, the proliferating cells and apoptotic cells were captured at room temperature using an OLYMPUS FLUOVIEW FV1000.

### Statistical analysis

The data were analysed with Novoexpress, ImageJ, statistical software in GraphPad Prism 8 for Windows (GraphPad Software). Quantitative data are presented as the means S.D. Experiments were performed at least three times for each experiment. NS, not significant, “*” p < 0.05, “**” p < 0.01, “***” p < 0.001 and “****” p < 0.0001.

## Results

### *Amd1* was highly expressed in hepatocytes during rapid liver growth

Early studies used microarrays to screen out some liver-enriched genes, and their roles in mouse and zebrafish liver development were proven (Cheng et al., 2006; Jochheim-Richter et al., 2006; Petkov et al., 2004), while many genes regulating liver growth were not identified using this method. Recently, bulk RNA-seq and sc-RNA-seq have been used for the study of liver regeneration and liver disease (Ben-Moshe et al., 2022; Saviano et al., 2020; Wang et al., 2019), but these methods have not been used to screen genes regulating liver growth. We found that during early zebrafish development, the liver undergoes rapid growth from 2dpf to 5dfp (Fig. 1 A, B) and hypothesized that genes regulating live growth should be highly expressed in hepatocytes during this stage. To identify genes being involved in liver growth, we performed RNA-seq using GFP-labelled hepatocytes sorted at different stages (Fig. 1C): 3dpf, 7dpf and adult zebrafish. The results showed that at 3dpf, 1654 genes were enriched in the hepatocytes, which was more than 2-fold higher than that in non-hepatocytes (Table S2). Among the top 25 genes of the 1654 genes, 16 genes had been reported to be liver-enriched genes in the Zfin database, 3 genes were reported in previously published literature, and 8 genes were not reported previously (Fig. 1 D and Table S3), implying that some new liver-enriched genes were identified. Among all 1654 liver-enriched genes, 262 genes were highly expressed in adult hepatocytes, which was 2-fold higher than that in 3dpf hepatocytes (Table. S4). Detailed analysis revealed that among the top 20 genes of the 262 genes, only 4 genes were not reported in early literature or in the zfin database (Table S5). In contrast, among the 1654 liver-enriched genes at 3dpf, 813 genes were highly expressed in 3dpf hepatocytes, which was more than 2-fold higher than that in adult hepatocytes (Table S6). To further confirm this result, 4 genes were randomly selected from the 813 genes, and their expression was evaluated in 3dpf hepatocytes and adult hepatocytes using RTLqPCR. The results showed that the expression of all of these genes was higher in 3dpf hepatocytes than in adult hepatocytes (Fig.1 E). These results implied that among the liver-enriched genes at 3dpf, the expression of most of them was higher in 3dpf hepatocytes than in adult hepatocytes, and they could be candidates for liver growth regulation.

**Figure 1.**
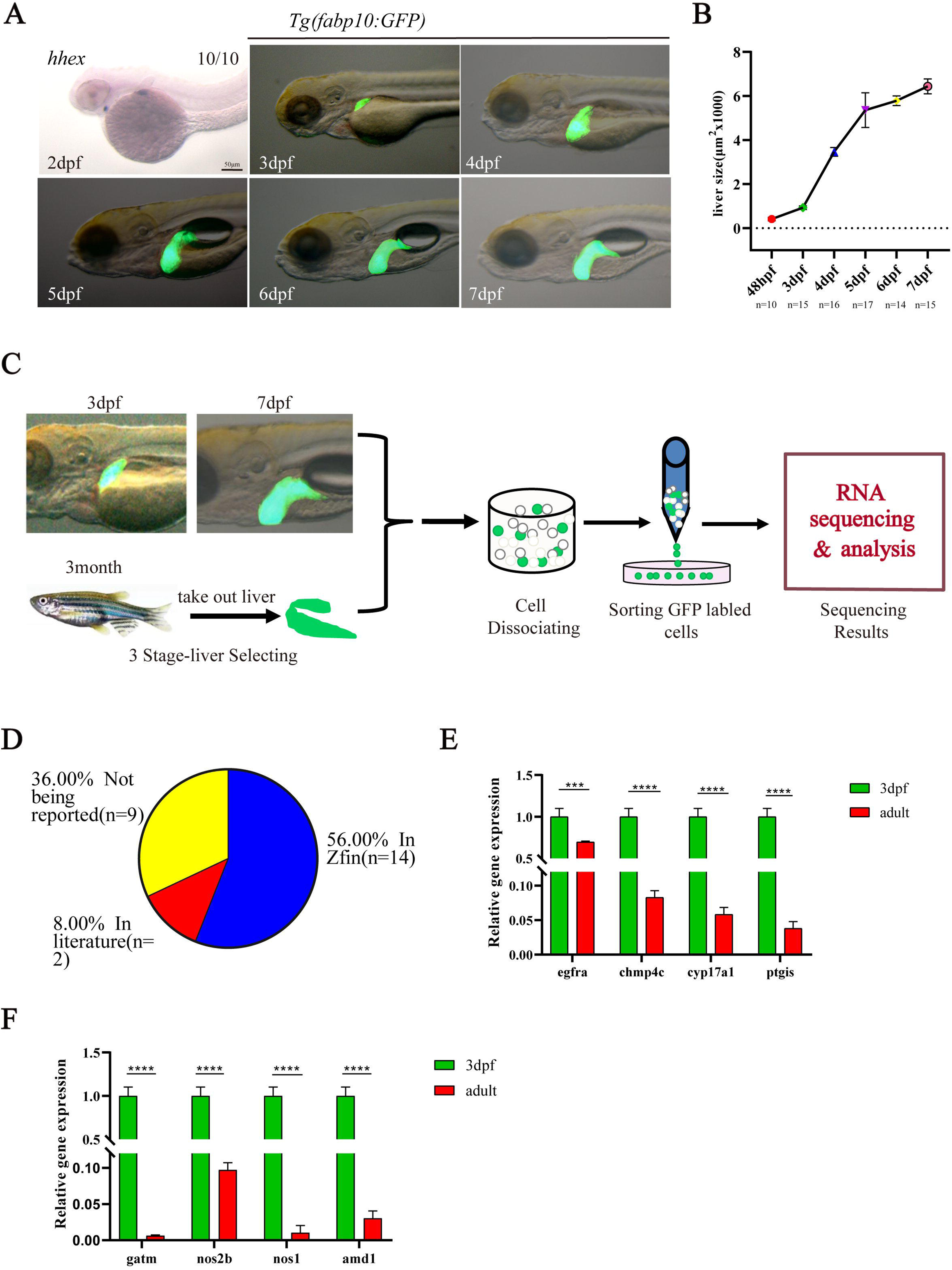
liver enriched gene screening and analysis. (A-B) Liver size comparing from 2dpf to 7dpf using *in situ* hybridization and living *Tg(fabp10:GFP)* transgenic embryos. Lateral view. (C) Schedule for sorting different staged-hepatocytes and bulk RNA sequencing (RNA-seq). (D) In the top 25 genes of liver-encriched genes on 3dpf, 56%, and 12% of them was reported to be enriched in embryonic liver in zfin database or being reported in early literatures. 32% of them is uncharacterized. (E) 4 genes were randomly selected from the genes being enriched and highly expressed in hepatocytes on 3dpf. The expression level of them was higher in hepatocytes on 3dpf than that in adult hepatocytes: the expression level in adult hepatocytes of *egfra,chmp4c*, *cyp17a1*and *ptgis* is 69.7%,8.3%, 5.8% and 3.8% of that in hepatocytes on 3dpf embryos, respectively. (F) The relative expression of *gatm*(0.63%), *nos2b*(9.7%), *nos1*(1.0%) and *amd1*(3.0%) in 3dpf-hepatocytes is much higher than that in adult-hepatocytes. Values are reported as mean ± SEM. “***” P < 0.001, “****” P < 0.0001. Scale bars, 50μm.

Since the ratio of liver growth was significantly decreased from 5dpf (Fig. 1A, B), we hypothesized that the expression of candidates regulating liver growth should be down regulated in hepatocytes at 7dpf compared with that in hepatocytes at 3dpf. Then, we analyzed the expression of 813 genes in hepatocytes at 3dpf and 7dpf. Indeed, the expression of 756 genes was decreased in hepatocytes at 7dpf (Table. S7). This result further indicated that the genes being identified here could be candidates for liver growth control. Next, we performed KEGG analysis to determine the signalling pathways associated with these genes. Interestingly, KEGG analysis showed that 4 genes related to arginine and proline metabolism were highly expressed in hepatocytes at 3dpf, and this result was confirmed by RTLqPCR experiments (Fig. 1F). Among these 4 genes, *nos1* has been reported to be involved in liver growth after injury (Cox et al., 2014); *amd1*, the key enzyme involved in the synthesis of polyamines (Pegg, 2009), has even been reported to be associated with ESC self-renewal and cell proliferation (James et al., 2018; Lim et al., 2018; Zhang et al., 2012; Zhao et al., 2012), but its role in liver development has not been addressed. We examined the role of *amd1* during liver growth to confirm that the genes we identified could be candidates for liver growth.

### The detailed expression pattern of *amd1* in early zebrafish development

To evaluate the possible role of *amd1* in liver growth, we first examined the expression of *amd1* in early zebrafish embryos using RTLPCR. The data showed that *amd1* was expressed in all staged embryos (Fig. 2A). To evaluate the detailed expression pattern of *amd1* during embryogenesis, we prepared *amd1* antisense/sense probes to determine its expression from 3 hours post fertilization (hpf) to 4dpf (Fig. 2B). The data showed that *amd1* was a maternal factor (Fig. 1 Bb1) and was expressed ubiquitously before 24 hpf (Fig. 2Bb1-b4). After 24 hpf, *amd1* was restricted to head and endodermal cells (Fig.2Bb5-b7). Importantly, *amd1* was highly expressed in the liver at 3 dpf and 4 dpf (Fig.2Bb6-8; Bb9-b11); this result was consistent with the RNA-seq data. In addition, detailed analysis of the liver growth ratio revealed that the liver undergoes the most rapid growth from 3 dpf to 4 dpf (Fig. 1 A, B), from 4 dpf, the ration of liver growth was decreased (Fig. 1B). Therefore, we hypothesized that the *amd1* expression level should be higher in hepatocyte in 3 dpf than that in 4 dpf. To evaluate this hypothesis, the relative expression level of *amd1* in liver hepatocytes at 3 dpf and 4 dpf was evaluated using sorted GFP-labelled hepatocytes at 3 dpf and 4 dpf (Fig. 2C, D). The data showed that the expression of *amd1* in hepatocytes in 3 dpf was higher than that in 4 dpf (Fig. 1. D). Here, the correlation between high expression of *amd1* and most rapid liver growth at 3 dpf strongly suggested the possible role of *amd1* in liver growth.

**Figure 2.**
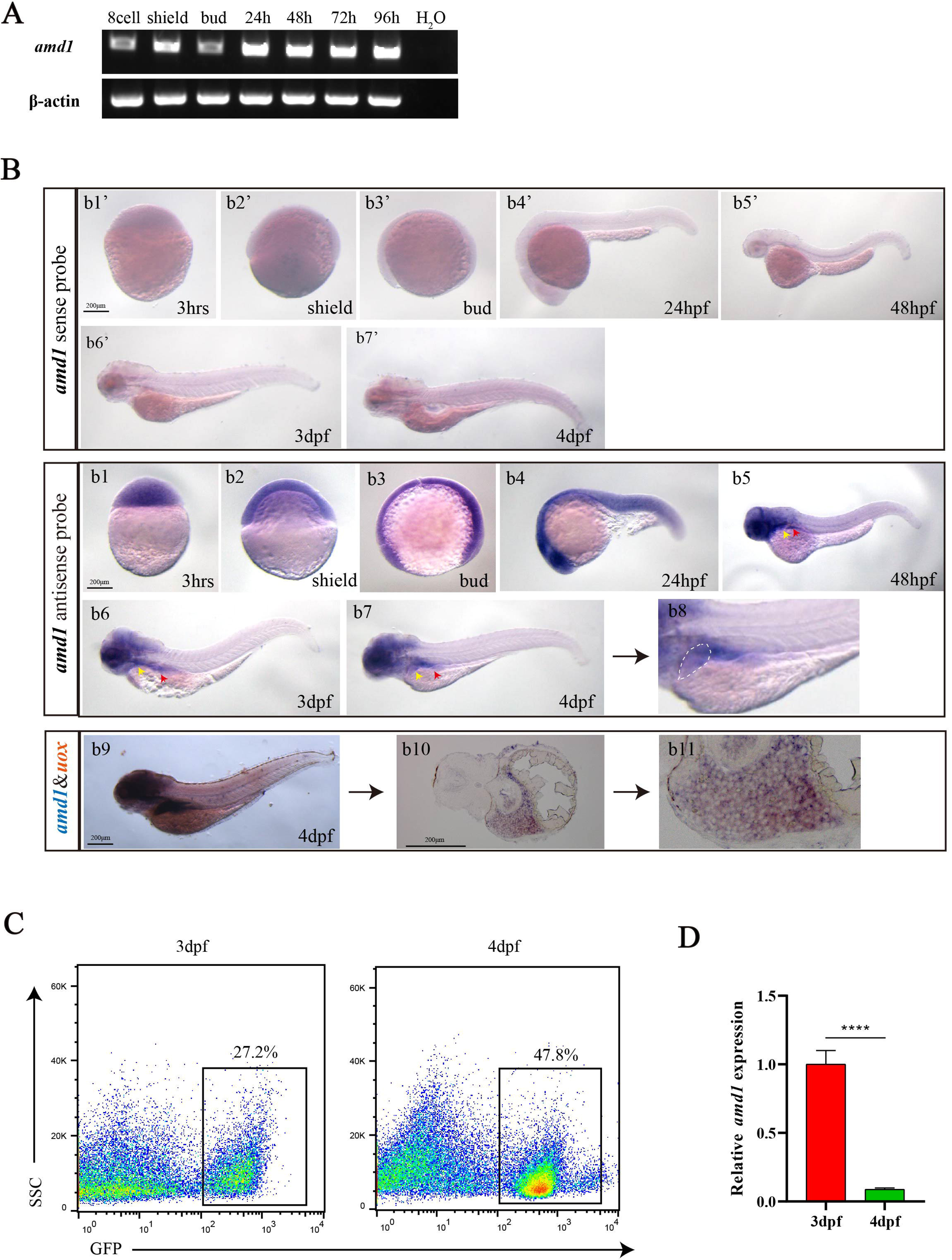
The expression pattern of *amd1* in early embryonic development. (A, B) The expression of *amd1* in early embryonic development. (A) PCR amplification for *amd1* in embryos at 8-cell stage, shield stage, bud stage, 24hpf, 48hpf, 72hpf and 96hpf. (B) *In situ* hybridization staining for *amd1* sense probe (Bb1’-b7’) and antisense probe (Bb1-b8). Especially from 2dpf to 4dpf, *amd1* was expressed highly in endoderm cells (Bb5-b8, red arrow showed), including in liver (Bb6-b8, yellow arrow and dashed box showed). (b10-b11) Double staining for *uox* (fast red) and *amd1*(blue) at 4dpf. *Uox* and *amd1* was colocalized in liver, amd1 was also expressed in gut. (C, D) Comparing the expression of *amd1* in hepatocytes on 3dpf and 4dpf using RT-qPCR. Sorting the GFP lableled hepatocytes on 3dpf and 4dpf (C). The data also showed the ratio of hepatocyte in embryos on 3dpf (27.2%) is largely lower than that on 4dpf (47.8%) (C). The expression level of *amd1* in hepatocytes on 4dpf is 8.9% of that in hepatocytes on 3dpf (D). Values are reported as mean ± SEM. “****” P < 0.0001. Scale bars, 200μm.

### *Amd1* mutant generation

To explore the role of *amd1* in zebrafish liver development, we used the CRISPRLCas9 method to construct the *amd1* mutant line as described in a previous report (Kroll et al., 2021). We designed one sgRNA for the target sequence that localized in exon 1 of *amd1* to construct the mutant lines (Fig. S1A). In F1 adult fishes, we screened two useful mutant lines: in mutant 1, the base group “CCCG” was changed to “TT”, and this exchange led to a prestop codon at 168 bp in the CD region (we named it *amd1^168^)*; in mutant 2, seven bases “CGGCAGG” were deleted, and a prestop codon arose at 141 bp in the CDs region (we named it *amd1^7^)* (Fig. S1B, C). Since the genome typing of *amd1^7-/-^* could be performed using PCR, we planed to use the *amd1^7-/-^* mutant line for most of the future experiments. Comparing the expression of *amd1* in mutants and wild-type embryos at 4 dpf suggested that the expression of *amd1* was decreased in *amd1^7-/-^* embryos (Fig. S1D). This result indicated that the premature mutant *amd1* mRNA was not stable and degraded during embryogenesis (Liu et al., 2019). Comparing the embryonic development between wild-type and *amd1^7-/-^* embryos, no distinct difference was discovered during early development (Fig. S1E, F), but the homozygotes were not viable and could not grow up to adult. This phenotype is similar to that in *Amd1 ^−/−^* mouse embryos, in which *Amd1 ^−/−^* mouse embryos died at the early developmental stage (Nishimura et al., 2002). All these results implied the critical role of *amd1* in embryonic development, and our *amd1* mutant lines could be used to address the role of *amd1* in liver development.

### Liver growth was repressed in *amd1* mutants

To observe the role of *amd1* in liver development easily, we crossed *amd1^7+/-^* with the *Tg(fabp10:GFP)* transgenic line to obtain *amd1* heterozygotes with the *Tg(fabp10:GFP)* transgenic background. Then, these fish were incrossed to obtain *amd1* homozygote embryos, and the liver phenotype was evaluated. The data showed that from 3 dpf to 5 dpf, the liver in *amd1^7-/-^* embryos was smaller than that in controls (Fig.3Aa1-a6, B). We also examined the liver size between *amd1* mutants and controls in non-transgenic embryos, the liver size was also smaller in *amd1^7-/-^* embryos at 3 dpf (Fig. 3C- F). Being similar, we also observed that the liver in *amd1^168-/-^*embryos was smaller than that in control (Fig. S2A-D). To further examine whether *amd1* was involved in liver specification during early embryogenesis, the early markers *prox1* and *hhex* were evaluated at 48 hpf (Shin et al., 2007). The data showed that the expression of *prox1* and *hhex* was intact in *amd1* mutants in the early stage (Fig. S3A, B). These results demonstrated that *amd1* loss of function only inhibited liver growth while not disturbed hepatocyte specification. Since *amd1* was also expressed in the gut at 3 dpf and 4 dpf (Fig. 2Bb7-b11), we examined whether gut growth was repressed in *amd1* mutants. The results showed that the gut marker *fabp2* (Parmar and Wright, 2013) was downregulated in *amd1* mutants (Fig. S4 A, B), implying that *amd1* also plays a critical role in gut development.

**Figure 3.**
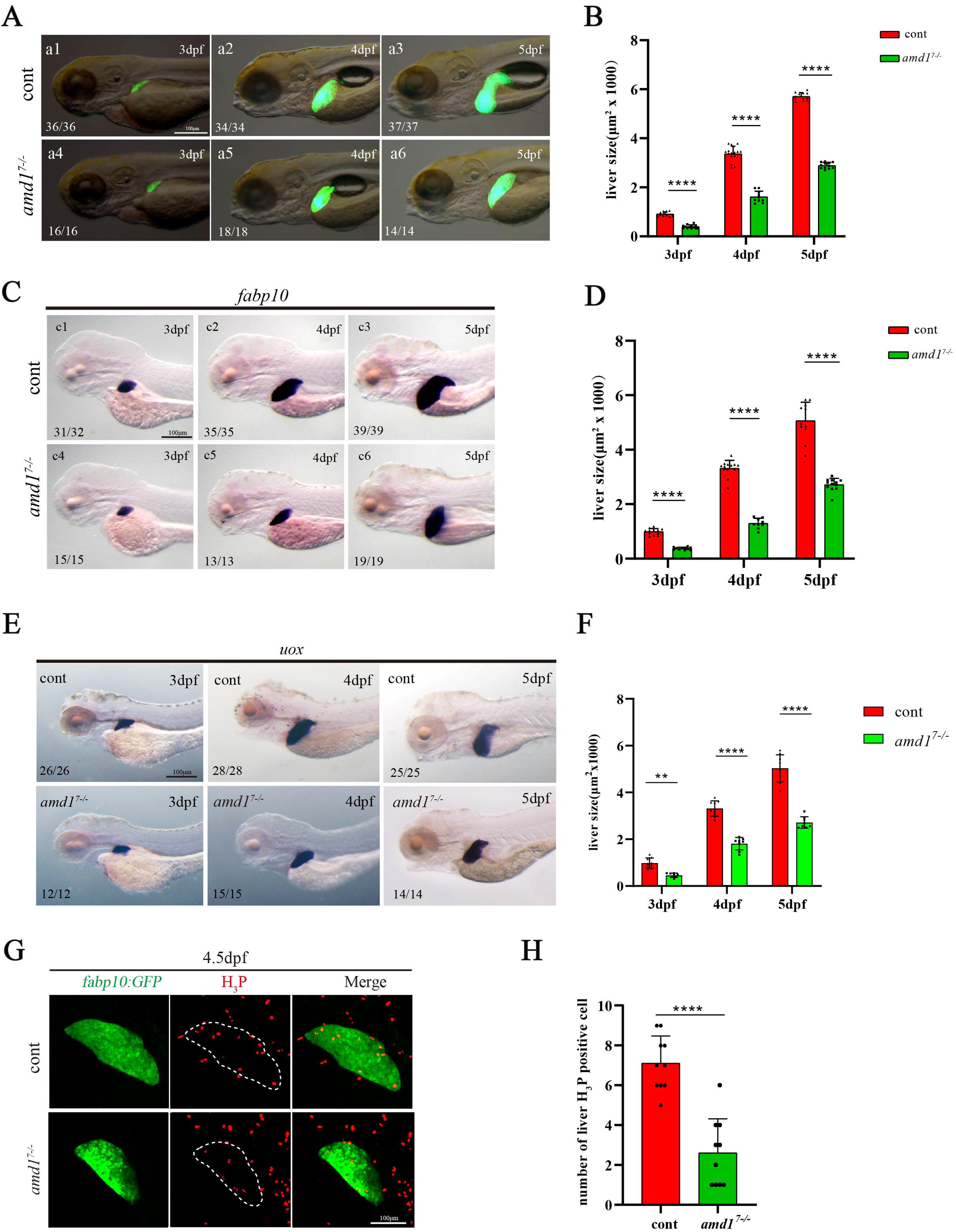
liver growth and hepatocyte proliferation were repressed in *amd1^7-/-^* liver. (A, B) Comparing the liver size in *amd1^7-/-^* embryos and control embryos using *Tg(fabp10:GFP)* transgenic line. From 2dpf to 5dpf, the liver size is smaller in *amd1^7-/-^* embryos (Aa1-a6). The size of liver in *amd1^7-/-^* embryos was 45.7% (n=16, p<0.0001), 52.2% (n=18, p<0.0001) and 49.3% (n=14, p<0.0001) of that in control embryos on 3dpf, 4dpf and 5dpf, respectively. (C, D) Comparing the liver size in *amd1^7-/-^* embryos and control embryos using *fabp10*staining (Cc1-c6). The size of liver in *amd1^7-/-^* embryos was 62.4% (n=15, p<0.0001), 61.0% (n=13, p<0.0001) and 46.3% (n=19, p<0.0001) of that in control embryos on 3dpf, 4dpf and 5dpf, respectively.(E, F) Comparing the liver size using *uox*staining (E). The size of liver in *amd1^7-/-^*embryos was 53.6% (n=12, p<0.01),45.5% (n=15, p<0.0001) and 46.0% (n=14, p<0.0001) of that in control embryos on 3dpf, 4dpf and 5dpf, respectively (F). (G, H) H_3_P staining for *amd1^7-/-^*embryos and control embryos. On 4.5dpf, the hepatocytes stainned with H3P in *amd1^7-/-^*embryos were decreased than that incontrol embryos G). In control embryos, approximate 7.1 hepatocytes were stained with H_3_P (n=10); in *amd1^7-/-^*embryos, approximate 2.6 hepatocyteswere stained with H_3_P (n=10, P< 0.0001) (B). Values are reported as mean ± SEM. NS, not significant, “****” P < 0.0001, Scale bars, 200μm.

### Hepatocyte proliferation was inhibited in *amd1* mutant embryos

During liver development, both hepatocyte proliferation delay and apoptosis increase were reported to give rise to a smaller liver (Chen et al., 2005; Chu and Sadler, 2009; Ma et al., 2019). To observe the underlying reason why the size of liver was reduced in *amd1* mutants, we examined hepatocyte proliferation and apoptosis in *amd1^7-/-^* embryos with a *Tg(fapb10a:GFP)* transgenic background. H3P immunostaining showed that hepatocyte proliferation was decreased in *amd1^7-/-^* embryos (Fig. 3G, H), but TUNEL experiment and Caspase3 immunostaining showed that the apoptosis of hepatocytes was not increased significantly in *amd1^7-/-^* embryos (Fig. S5B-E). To further confirm the role of *amd1* in regulating proliferation, the proliferating related markers were examined using RT-qPCR and the data demonstrated that the expression of *cdk1*, *cdk4*, *chk1* and *mcm5* was downregulated in *amd1* mutants (Fig. S5A). This result above further demonstrated that *amd1* is required for hepatocyte proliferation during liver growth, which is consistent with that being reported in early literature, in which AMD1 is required for EST self-renewal and cell proliferation (James et al., 2018; Zhang et al., 2012).

### *Skp2* expression was downregulated in *amd1^7-/-^* embryos

Early research showed that AMD1 played multiple roles during normal embryo development and disease, including the regulation of polyamine synthesis and gene transcription (Bian et al., 2021; James et al., 2018; Patel et al., 2018; Zabala-Letona et al., 2017). Mechanistically, AMD was reported to lie downstream of c-Myc, C/EBP/β and mTORC1 (Snezhkina et al., 2016; Zabala-Letona et al., 2017), and it also lies upstream of MINDY1(James et al., 2018). Therefore, the mechanism by which *amd1* regulates hepatocyte proliferation during liver growth is complicated. To elucidate the detailed role of *amd1* in liver growth, we analysed the gene expression in *amd1^7-/-^* embryos at 4 dpf using RNA-seq. The results showed that in *amd1^7-/-^* embryos, 94 genes were upregulated, and 104 genes were downregulated (Fig. 4A). Among the downregulated genes, KEGG analysis showed that four genes were in the mTOR signalling pathway, including *skp2* (Fig. 4B C). Detailed evaluation using RTLqPCR revealed that both *skp2* and the *skp2* downstream gene *rho* were downregulated (Fig. 4D). *In situ* experiments further showed that *skp2* was also downregulated in *amd1^7-/-^* embryos on 3 dpf (Fig. 4E), especially in the liver and gut (Fig. 4 E). In addition, we further examined the detailed expression pattern of *skp2* in early embryonic development using *in situ* experiments. The data showed *skp2* was a maternally expressed gene (Fig. 4Ff1, f2), it was expressed ubiquitously before 24 hpf (Fig. 4Ff3- f5) and then restricted to the liver, gut, eyes and boundary of the hindbrain and midbrain at 3 dpf (Fig. 4Ff6). The enriched expression of *skp2* in liver and gut at 3 dpf further implied the role of *skp2* in liver growth. In conclusion, all the results above suggested the possibility that *skp2* is required for *amd1* to regulate liver growth during embryonic development.

**Figure 4.**
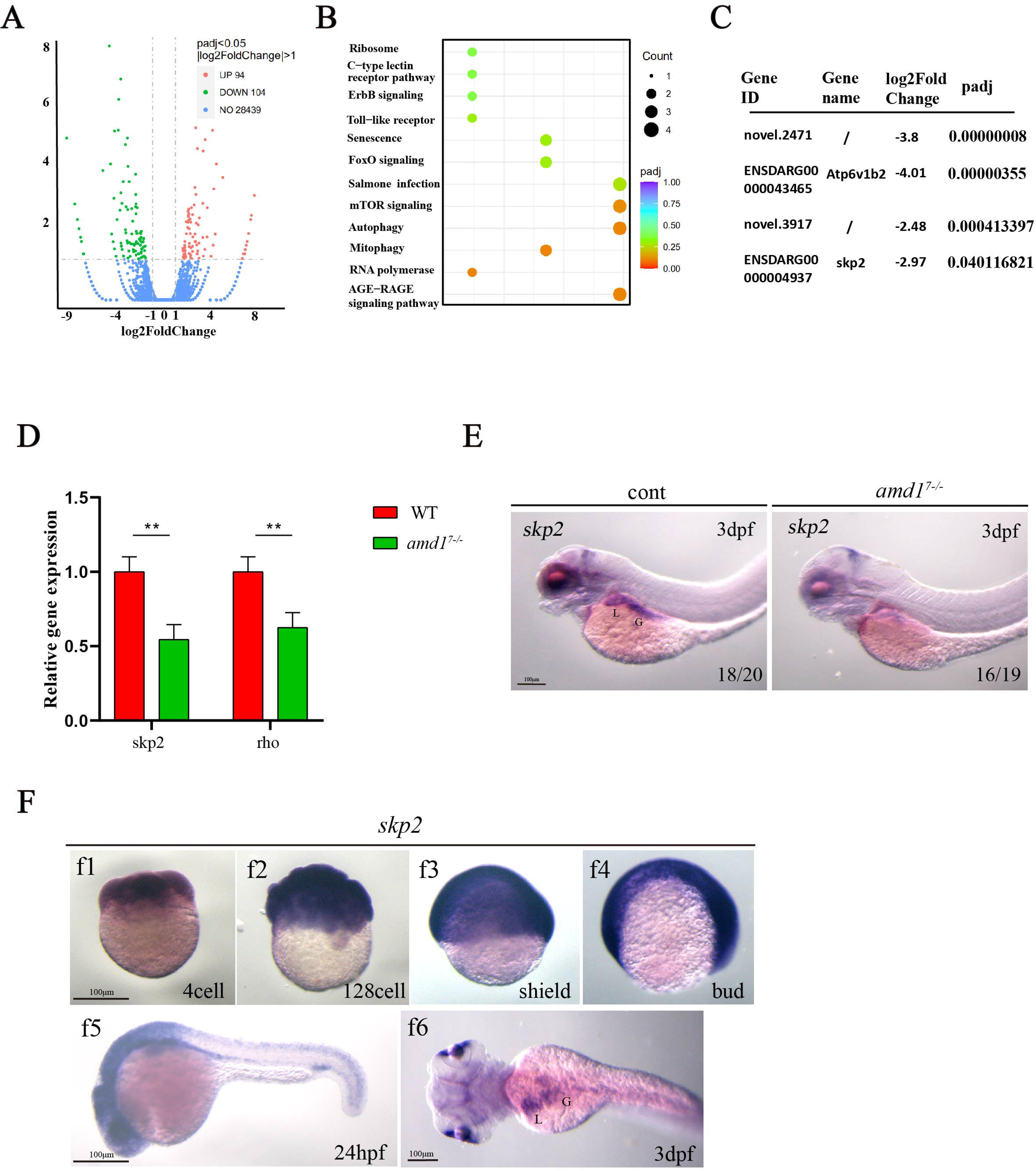
*skp2* was downregulated in *amd1^7-/-^*embryos. (A-C) RNA sequencing data analyisis. In *amd1^7-/-^* embryos, 94 genes were up-regulated, 104 genes were down-regulated, there is no significant difference for the expresion of 28439 genes between cotrols and *amd1^7-/-^* embryos (A). In the down-regulated genes, some of them were belonged to mTOR signaling (B). *skp2* was significantly downregulated in *amd1^7-/-^* embryos (C). (D) RT-qPCR experiments showed that *skp2* (0.545 folds to control, p=0.0011) and *skp2* downstream gene *rho* (0.626 folds to control, p=0.0036) was downregulated in *amd1^7-/-^*embryos. (E) *In situ* experiments showed that *skp2* was downregulated in liver and gut in *amd1^7-/-^*embryos on 4dpf. (F) The expression of *skp2* was examined at 4-cell stage (f1), 128-cell stage (f2), shield stage (f3), bud stage (f4), 24hpf (f5) and 3dpf (f6). On 3dpf, *skp2* was enriched in eyes, pancreas, liver and gut (f6). Values are reported as mean ± SEM. “**” P < 0.01. Scale bars,100μm.

**Figure 5.**
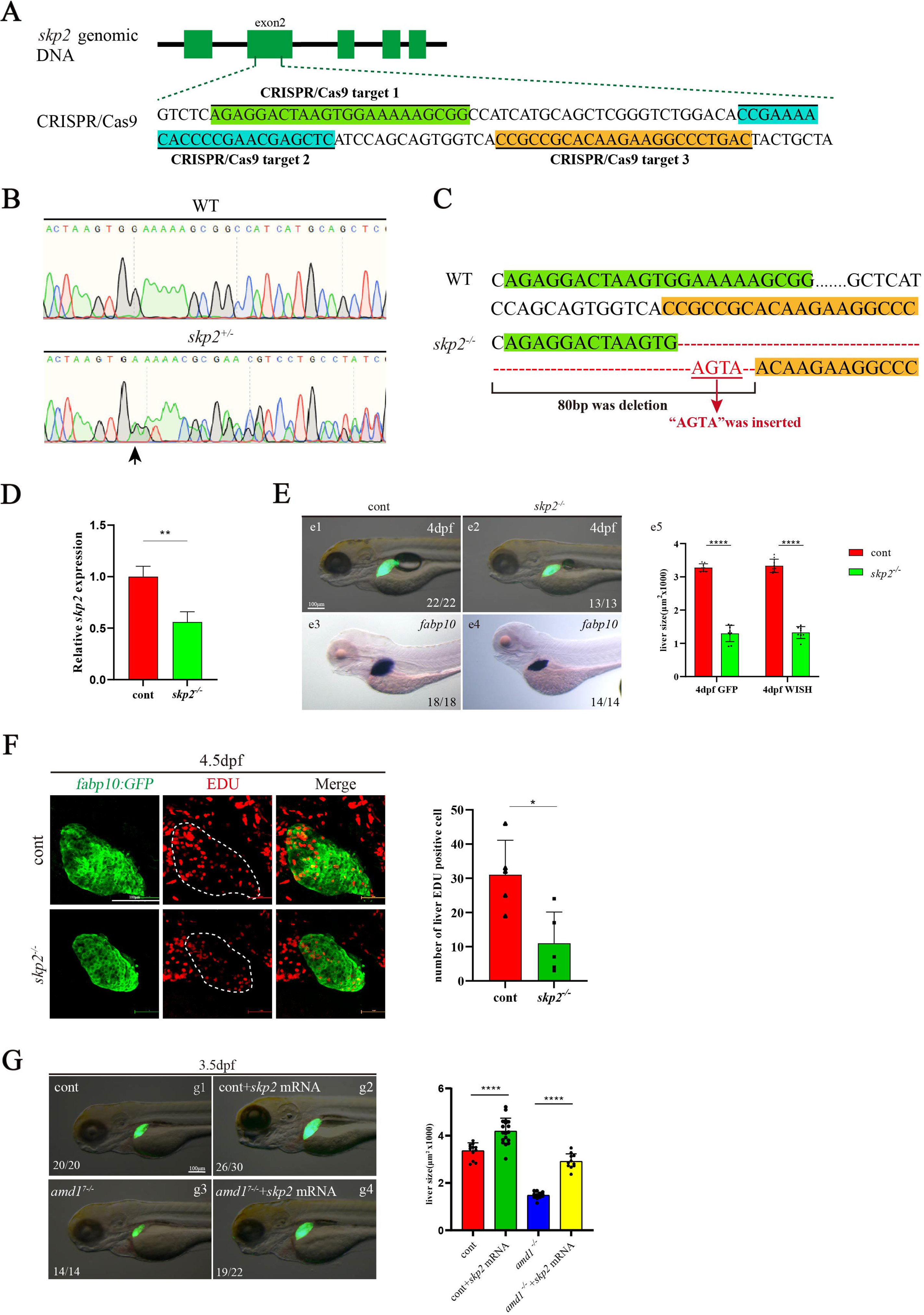
*skp2* is required for liver development and lies downstream of *amd1*. (A) Three targets of *skp2* sgRNA in exon2 of *skp2* gene. (B) The sequencing results of *skp2* wild type (up sequence) and *skp2^+/-^* embryos (down sequence). (C) 80 nucleotides between “AAAAAGCGG” (green line labelled) and “ACAAGAAGG” (yellow line labelled) were deleted, but “AGTA” were added in these two sequences. This mutantion gave rise to a frame-shift and there is no stop codon at the terminal region of *skp2* CDs. (D) The expression of *skp2* in *skp2^-/-^* mutants was 55.2% of that in wild type controls on 4dpf. Values are reported as mean ± SEM. “**” P < 0.01. (E) Liver size was compared in controls and *skp2* mutants using *Tg(fabp10:GFP)* transgenic line and wild type line embryos. The liver size in *skp2* mutants (e2, e4) is smaller that in controls (e1, e3). Statistical analysis was did for the liver size in *skp2* mutants and controls (e5). Values are reported as mean ± SEM. “****” P < 0.0001. (F) Cell proliferation was examined using Edu staining on 4.5dpf. The number of hepatocytes staining with Edu in *skp2* mutants is smaller than that in controls. Values are reported as mean ± SEM. “**” P < 0.01. (G) *skp2* overexpression increased the size of liver. On 3.5 dpf, in wild type embryos injection of *skp2* mRNA increased the size of liver (86.6%, n=18); meanwhile the phenotype “smaller liver” in *amd1^7-/-^*embryos was rescued in 86.3% of embryos by injecting *skp2* mRNA (n=11). “*” P < 0.05, “***” P < 0.001, “****” P < 0.0001, Scale bars, 100μm.

### *Skp2* plays a critical role in liver development

Skp2, a key component of the SKP1-cullin 1-F-box (SCF) complex (Wu et al., 2021; Zhang et al., 1995), largely functions as an oncoprotein (Cai et al., 2020). Previous work showed that Skp2 is involved in cell proliferation, migration, invasion and metastasis in some malignant tumours (Cai et al., 2020; Wang et al., 2012). In HCC, Skp2 was also involved in HCC metastasis (Chen et al., 2021; Zhang et al., 2017).These studies above implied the possibility that *skp2* is essential for *amd1* to regulate liver development in zebrafish. To evaluate this hypothesis, first we generated *skp2* mosaic mutation embryos using the CRISPR/Cas9 method (Fig. 5A and Fig. S6A)(Kroll et al., 2021) and examined liver development in *Tg(fabp10:GFP)* transgenic lines. After the exon2 of *skp2* was edited in mosaic mutants (Fig. S6A-C), the early embryonic development was delayed (Fig. S6D) and liver growth (Fig. S6E) was repressed. Next we generated the stable *skp2* mutant line to confirm the role of *skp2* in liver growth. We screened out a frameshift mutant line for *skp2*: in this mutant 80 nucleotides between “AAAAAGCGG” (labeled with green) and “ACAAGAAGG” (labeled with yellow) were deleted, but “AGTA” was inserted between these two sequences (Fig. 5A-C). This mutantion gave rise to a frameshift and there is no stop codon at the terminal region of *skp2* CDs. RT-qPCR further showed that the level of *skp2* was decreased in *skp2* mutant embryos at 3 dpf (Fig. 5D). Further, we found that the *skp2* homozygotes did not display distinct external phenotype at early developmental stage (Fig. S7) but were not viable and could not grow up to adulthood. The data also showed that the liver size was decreased (Fig. 5Ee2, e4, e5 and Fig. S8A-D), as well the hepatocytes proliferation was decreased (Fig. 5F), demonstrating the critical role of *skp2* in liver growth. Finally, we examined whether over-expressing *skp2* by injection of *skp2* mRNA could restore the liver development in *amd1* mutant. The data showed that, overexpression of *skp2* partially increased the liver size in control embryos (Fig. 5Gg1, g2) and restored the liver size in *amd1* mutants (Fig. 5Gg3, g4). In conclusion, all these data suggested that *skp2* is required liver development, it is also essential for *amd1* to regulate liver development.

### *Skp2* is required for *amd1* to regulated liver development at liver rapid growth stage

Since *sk*p2 was ubiquitously expressed at early stage, we further evaluated whether *skp2* is also involved in early liver development. The data showed that the expression of *prox1* and *hhex* was decreased in *skp2* mutants at 48 hpf (Fig. S9A, B), implying *skp2* is also involved in early liver development. This result gave rise to a possibility that the liver phenotype is a secondary result of early developmental defects in *skp2* mutants. To observe the direct role of *skp2* at liver growth stage, we used the *skp2* inhibitor SMIP004 (Li et al., 2019) to block the activity of *skp2* when the liver undergoes rapid growth (Fig. 6A). During concentration titration, we found that treatment with 40 µM SMIP004 did not lead to gastrulation defects or smaller livers (Fig. 6A-C), while treatment with 80 µM SMIP004 decreased the activity of Skp2, displaying decreased expression of *rho* and *ahcy* (Fig. 6 E), the two downstream genes of *skp2* (Cai et al., 2020). Then, we used 80 µM SMIP004 to treat embryos from 2 dpf to 4 dpf and analysed liver size (Fig. 6A, F). The data showed that *skp2* inhibition did not lead to distinc embryonic phenotype (Fig. 6D) but resulted in smaller liver at 4 dpf in *Tg(fabp10:GFP)* transgenic embryos and wild-type embryos (Fig. 6Gg2, Hh2). This result confirmed that *skp2* played a vital role during liver growth. In addition, *skp2* inhibition decreased hepatocyte proliferation (Fig.6Ii2, i5), confirming the role of *skp2* in liver growth. To further evaluate whether *skp2* is required for *amd1* to regulate liver growth, we determined whether treatment with SMIP004 gave rise to a much smaller liver and much less proliferating hepatocyte in *amd1^7-/-^* embryos. The data showed that SMIP004 treatment further reduced the size of the liver in *amd1^7-/-^* embryos (Fig. 6Gg4, Hh4), and the H3P labelled cells was decreased (Fig. 6Ii8). In conclusion, all the data above showed that *skp2* is reqiured for *amd1* to regulate liver growth at liver rapid growth stage.

**Figure 6.**
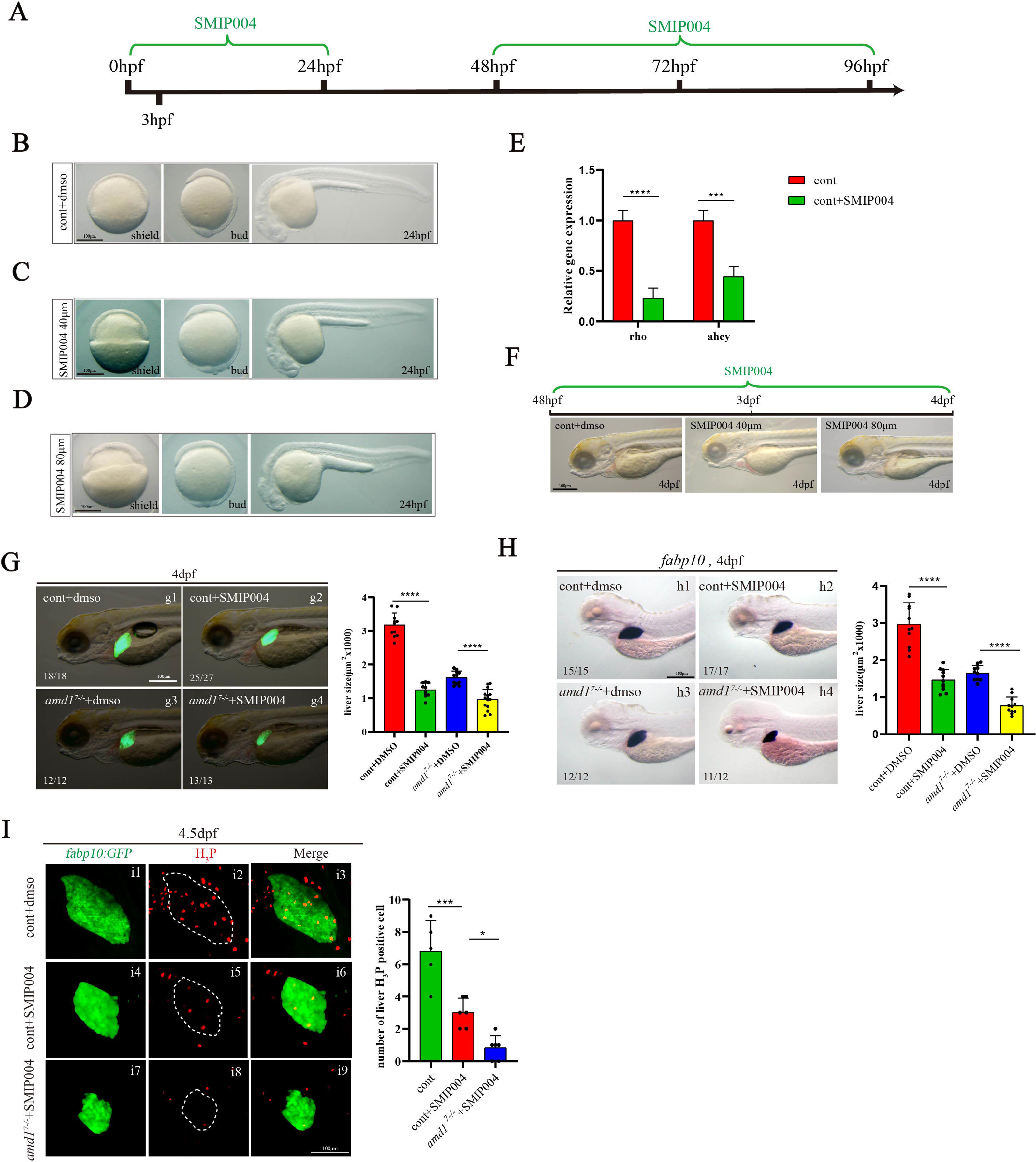
*skp2* is required for liver growth and mediates *amd1* regulates liver development. (A) The live phenotype after downregulatio of *skp2* using mosaic knock out on 3.5dpf.Most of embryos (88.1%, n=12, p< 0.0001) displayed smaller liver in embryos injected with Cas9/*skp2* sgRNAs.(B)Schedule for inhibiting *skp2* activity using SMIP004 treatment. (C) After inhibiting *skp2* activity, 92.6% of embryos (Cc2, n=10, p< 0.0001) displayed smaller liver than that in controls (Cc1, n=11); in *amd1^7-/-^*embryos, skp2 inhibition (Cc4, 100%, n=13, p< 0.0001) made the liver much smaller that controls (Cc3, n=12). (D) *In situ* experiment also showed that, 100% of embryos (Dd2, n=11) displayed smaller liver than that in controls (Dd1, n=11). Inhibiting *skp2* activity made most of *amd1^7-/-^* embryos (Dd4, 91.6%, n=11, p< 0.0001) displayed much smaller liver that controls (Dd3, n=10). (E) Cell proliferation evaluation using H_3_P staining. After treatment with SMIP004, H_3_P staining hepatocytes were decreased comparing with control (Control, n=5; SMIP treatment, n=6, p=0.0004). SMIP004 treatment further decreased the number of hepatocytes staining with H3p (n=6, p=0.171). (F) *skp2* overexpression increased the size of liver. On 3.5 dpf, in wild type embryos injection of *skp2* mRNA increased the size of liver (86.6%, n=18); meanwhile the phenotype smaller liver in *amd1^7-/-^* embryos was rescued in 86.3% of embryos by injecting *skp2* mRNA (n=11, p=< 0.0001). Values are reported as mean ± SEM. “*” P < 0.05, “***” P < 0.001, “****” P < 0.0001, Scale bars,100μm.

### *Amd1-skp2* cascade plays a critical role during hepatocyte proliferation in a zebrafish HCC model

Both hepatocytes in developing liver and HCC were characterized by rapid cell proliferation (Chaturantabut et al., 2019; Perugorria et al., 2019). The role of *amd1* in regulating liver growth implied that *amd1* plays a vital role during hepatocyte proliferation in zebrafish HCC progression. To evaluate this hypothesis, we generated an HCC model in zebrafish larvae as described in previous report ((Yang et al., 2019) and Fig. S10A). In the HCC model, liver-specific overexpression of *Kras^G12V^* gave rise to a much larger liver compared with control embryos (Fig. S10Bb3, b6), and the expression of 5 HCC markers was greatly upregulated (Fig. S10C), indicating the HCC model was generated successfully. Next, we sorted GFP-labelled hepatocytes from normal liver and HCC model, then analysed the expression of *amd1* (Fig. 7A, B). The data showed that *amd1* was upregulated in HCC cells (Fig.7B). Further data showed that *amd1* loss of function decreased the size of liver in the HCC model (Fig. 7Cc3, D), and hepatocyte proliferation was also decreased (Fig. 7Ee5, F). These results demonstrated the role of *amd1* in hepatocyte proliferation was conserved in zebrafish HCC model.

**Figure 7.**
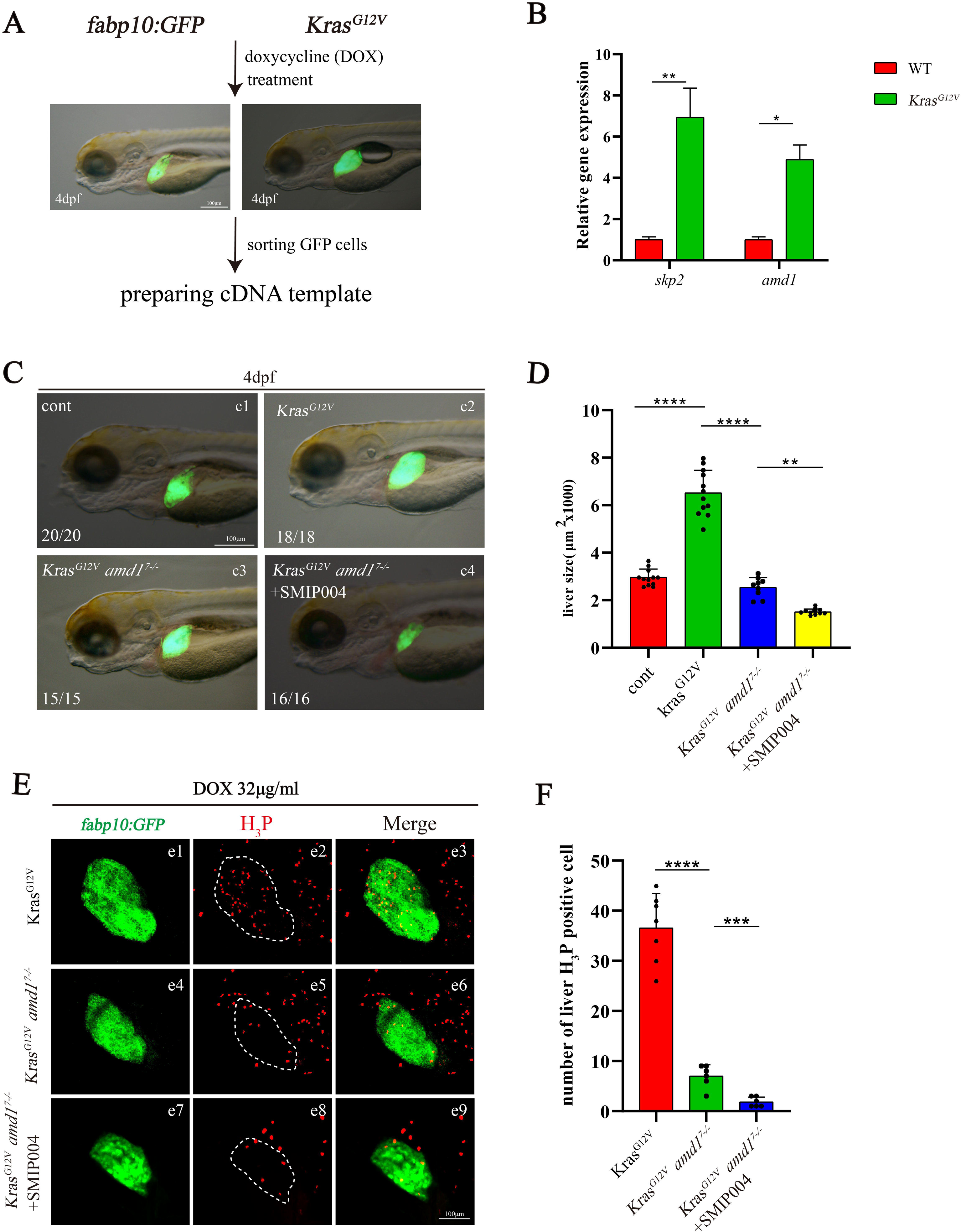
*amd1-skp2* cascade is reqiured for hepatocyte proliferation in zebrafish HCC model. (A) The schedule for examing the expression of *amd1* and *skp2* in hepatocytes for normal liver and HCC model. (B) Comparing with that in normal liver, the expression of *amd1* (5.39 folds to normal liver, p=0.0162) and *skp2* (7.93 folds to normal liver, p=0.0035) was increased significantly. (C, D) dox induced overexpression of *Kras* increased the liver growth (Cc2, n=18; D, n=12, p=< 0.0001), amd1 loss of function decreased the size of liver in HCC model (Cc3, n=15; D, n=9, p=< 0.0001), simultaniously inhibiting *skp2* and *amd1* made the liver much smaller in HCC model (Cc4, n=16; D, n=11, p=0.0012). (E, F) proliferating hepatocytes staining with H3p. Comparing with control, *amd1* loss of function decreased the number of H3p staining hepatocytes (Ee5; F, n=6, p=< 0.0001), simultaniously inhibiting *skp2* and *amd1* furtherdecreased the number of H3p staining hepatocytes (Ee8; F, n=6, p=0.0005). Values are reported as mean ± SEM. “*” P < 0.05, “**” P < 0.01, “***” P < 0.001, “****” P < 0.0001, Scale bars,100μm.

To evaluate whether *skp2* mediates *amd1* to regulate hepatocyte proliferation in the zebrafish HCC model, first the expression of *skp2* was examined. The data showed that the expression of *skp2* was also significantly upregulated in hepatocytes in the zebrafish HCC model (Fig.7B). Next, we determined whether *skp2* was required for zebrafish HCC and whether the liver size was much smaller in *amd1^7-/-^* embryos after blocking *skp2* activity. Indeed, the size of the liver was much smaller in *amd1^7-/-^* embryos treated with SMIP004 from 48hpf to 4dpf in zebrafish HCC model (Fig. 7Cc4, D), and hepatocyte proliferation was also decreased in *amd1^7-/-^* embryos treated with SMIP004 (Fig. 7Ee8, F). These results demonstrated the possibility that *skp2* also is required for *amd1* to regulate hepatocyte proliferation in a zebrafish HCC model.

## Discussion

Hepatocyte proliferation is one of the critical events during liver organogenesis and HCC progression. Even though many genes have been reported to be involved in this process (Cox et al., 2016; Cox et al., 2018; Wu et al., 2022; Yang et al., 2019), the underlying mechanism is far from being elucidated. Here, we screened 813 genes that were highly expressed in hepatocytes at 3 dpf. Meanwhile, we selected one of them, *amd1*, to evaluate whether it plays a crucial role during normal liver growth and HCC. We identified the role of *amd1* in hepatocyte proliferation during normal liver development and HCC progression and proved that *skp2* mediated, at least partially, *amd1* to regulate liver growth during liver development and HCC progression.

For the genes highly expressed in hepatocytes at 3 dpf, some of them, such as *Lats1*, *Rhbg* and *Npas2*, have been reported to be associated with liver growth or hepatocyte survival (Leibing et al., 2018; Yi et al., 2016; Yuan et al., 2017), demonstrating that the genes identified in our research could be the subjects for further studies. Noticeably, in our screening work, some genes were highly expressed in hepatocytes at 3 dpf but not specifically enriched in hepatocytes, possibly these kinds of genes could also be candidates for liver growth regulation. To this point, the evidence was that *hdac3, Id2a* and *oestrogen* have been reported to be involved in liver growth (Chaturantabut et al., 2020; Farooq et al., 2008; Khaliq et al., 2015)[2, 10,], but they are not specifically enriched in hepatocytes at 3 dpf. Of course, our RNA-seq work did not identify all the genes regulating liver growth. For example, Yap has been reported to regulate liver growth (Cox et al., 2016; Cox et al., 2018), but it was not found in our gene list. Therefore, more work is needed to further evaluate the role of the genes being identified in liver growth control.

*Amd1*, one of the genes we identified, was previously reported to be associated with ESC self-renewal, cell proliferation and cell migration (James et al., 2018; Lim et al., 2018; Zhang et al., 2012; Zhao et al., 2012), while its detailed role in liver development was not addressed. Our current data demonstrated that it is crucial for liver growth. When investigating the mechanism underlying, we identified *skp2* as one of the key downstream genes that regulate hepatocyte proliferation during liver growth. Even in early mouse embryos, Myc functions as the critical mediator for Amd1 to control ESC self-renewal (Zhang et al., 2012; Zhao et al., 2012), while during zebrafish liver outgrowth, we did not find that the expression of *myc* was significantly downregulated in *amd1* mutants. In contrast, *skp2* seems to mediate the role of *amd1* in regulating hepatocyte proliferation. These results are consistent with early reports in mice, in which c-Myc was reported to be dispensable for normal liver growth during the postnatal period (Baena et al., 2005; Sanders et al., 2012). In addition to the role of *amd1* in liver growth in zebrafish development, in a previous study, Amd1 was reported to stabilize the interaction of IQGAP1 with FTO, which upregulating the expression of *nanog*, *kif4* and *sox2* to accelerate HCC progression (Bian et al., 2021). Here, we further identified that the Amd1-Skp2 cascade played a vital role in hepatocyte proliferation in a zebrafish HCC model. In our zebrafish HCC model, the RNA-seq data and RTLqPCR showed that both *amd1* and *skp2* were significantly increased (Fig. 7B), and the liver size was reduced when the role of *amd1* or *skp2* was blocked (Fig. 7C-F). These data implied that the mechanism by which *amd1* regulates hepatocyte proliferation is complicated.

In conclusion, our work identified some uncharacterized genes enriched in hepatocytes at 3 dpf. These genes may be candidates for regulating hepatocyte proliferation in normal liver development and HCC. As one of these genes, *amd1* was identified to play a crucial role in regulating hepatocyte proliferation via *skp2* in normal liver outgrowth and HCC progression.

## List of abbreviations

HCC: hepatocellular carcinoma
AMD1: S-adenosylmethionine decarboxylase proenzyme
BMPs: bone morphogenetic proteins
RNA-seq: bulk RNA sequencing
scRNA-seq: single-cell RNA sequencing
ESC: embryonic stem cell
skp2: S-phase kinase-associated protein 2
RNP: ribonucleoprotein complex
hpf: hours post fertilization
dpf: days post fertilization.

## Informed Consent Statement

Not applicable.

## Data Availability Statement

All the data was in the manuscript and supplementary materials.

## Acknowledgements

We would like to thank Dr Chi Liu to help sorting GFP labelled hepatocytes, Xiaojun Yang to read and gave critical comments. We also would like to thank the members working in our fish facility to help take care of all the fish lines in this study.

## Author contributions

Conceptualization: SZH, ZHG; Methodology: SZH, KZ, ZHG; Investigation: KZ, XYW, BYC, YD, YL; Visualization: SZH, KZ, YD, ZHG; Funding acquisition: SZH;Project administration: SZH; Supervision: SZH; Writing – original draft: SZH, ZHG, KZ,ML; Writing – review & editing: SZH, ZHG, XDL, ML.

